# Inferring macro-ecological patterns from local species’ occurrences

**DOI:** 10.1101/387456

**Authors:** Anna Tovo, Marco Formentin, Samir Suweis, Samuele Stivanello, Sandro Azaele, Amos Maritan

## Abstract

1. Biodiversity provides support for life, vital provisions, regulating services and has positive cultural impacts. It is therefore important to have accurate methods to measure biodiversity, in order to safeguard it when we discover it to be threatened. For practical reasons, biodiversity is usually measured at fine scales whereas diversity issues (e.g. conservation) interest regional or global scales. Moreover, biodiversity may change across spatial scales. It is therefore a key challenge to be able to translate local information on biodiversity into global patterns.
2. Many databases give no information about the abundances of a species within an area, but only its occurrence in each of the surveyed plots. In this paper, we introduce an analytical framework to infer species richness and abundances at large spatial scales in biodiversity-rich ecosystems when species presence/absence information is available on various scattered samples (i.e. upscaling).
3. This framework is based on the scale-invariance property of the negative binomial. Our approach allows to infer and link within a unique framework important and well-known biodiversity patterns of ecological theory, such as the Species Accumulation Curve (SAC) and the Relative Species Abundance (RSA) as well as a new emergent pattern, which is the Relative Species Occupancy (RSO).
4. Our estimates are robust and accurate, as confirmed by tests performed on both in silico-generated and real forests. We demonstrate the accuracy of our predictions using data from two well-studied forest stands. Moreover, we compared our results with other popular methods proposed in the literature to infer species richness from presence-absence data and we showed that our framework gives better estimates. It has thus important applications to biodiversity research and conservation practice.

## Introduction

The problem of inferring total biodiversity when only scattered samples are observed is a long-standing problem. In the 1940s, the British chemist and naturalist A. S. Corbet spent two years in Malaya to trap butterflies [1]. For every species he saw, he noted down how many individuals of that species he trapped. When Corbet returned to England, he showed the table to its colleague R. A. Fisher and asked him how many new species he would trap if he returned to Malaya for another couple of years. The father of statistics was only the first to tackle the problem of species estimation [2], which since then has found large applications in different scientific fields, from ecology [3, 4, 5] to bioscience [6, 7, 8], leading to the development of a myriad of estimators [9, 10, 11].

Indeed, although ecological drivers crucial for conservations act at large scales, biodiversity is typically monitored at limited spatial scales [12, 13]. Extrapolating species richness from the local to the whole ecosystem scale is not straightforward, because it is not additive as a function of the area. As a result, a huge number of biodiversity estimators have been proposed in ecological literature [4, 14, 15, 16, 17, 18, 19, 20, 21]. Their commonest limitation is to have a limited application range (local/regional-scale extrapolations), and to be sensitive to the trees’ spatial distribution [22, 23, 24], sample coverage and sampling methods [25].

Many analytical methods have been proposed to upscale species richness using as input the local Relative Species Abundance distribution (RSA) [26, 27, 28, 29], i.e. the list of the species present at the sampled scale along with the proportion of individuals belonging to each of them. For example, estimates of biodiversity at large scales have been performed using log-series as the RSA [2]. The log-series distribution is often used to describe RSA patterns in many different ecological communities, characterised by high biodiversity [24]. Thanks to the availability and reliability of the species abundance data in forests (given by systematic and periodic field campaigns and high detectability of species), this method has been typically applied to tropical forests. In particular, it has been used to estimate the species richness of the Amazonia [27] and the global tropical tree species richness [28].

These methods have been proved to typically perform better than non-parametric estimators of biodiversity [30]. In contrast with the former, non-parametric approaches do not assume a specific family of probability distributions. In particular, non-parametric methods do not make any assumption on the RSA distribution and they thus perform no fit of empirical patterns, rather they only take into account rare species, which are intuitively assumed to carry all the needed information on the undetected species in a sample.

Nevertheless, all the aforementioned methods need abundance data in order to infer biodiversity at larger scale. However, in many open-access databases (e.g. species abundance data obtained from metagenomics [31]) this information is highly imprecise, if available at all. Indeed, there are lots of datasets which give only information about the presence or absence of a species in different surveyed plots, without specifying the number of individuals within them. Some non-parametric approaches have been generalized to infer species richness from this presence-absence data [30, 11]. Table 2 summarizes the most popular estimators and for each one details the predicted biodiversity as a function of the input data. However, most of them have the strong limitation that they do not have an explicit dependence of the observation scale, leading to poor estimates of the number of species at the global scales (see Results). The only estimator which takes into account the ratio between the surveyed area and the global one is the one introduced by Chao [30, 25, 11] and denoted here as *Chao*_wor_ (see Table 2). This method takes into account the number of species detected in one sample only and those detected in exactly two samples observed at the sample scale to infer the total species richness at the whole forest scale. However, it has been shown that Chao’s method, although giving reliable species estimates, it does not properly capture the empirical Species Accumulation Curve (SAC) [29], which describes how the number of species changes across spatial scales. In absence of spatial correlation, it is equivalent to another macro-ecological pattern of interest which is the Species Area Relation (SAR).

**Table 2:**
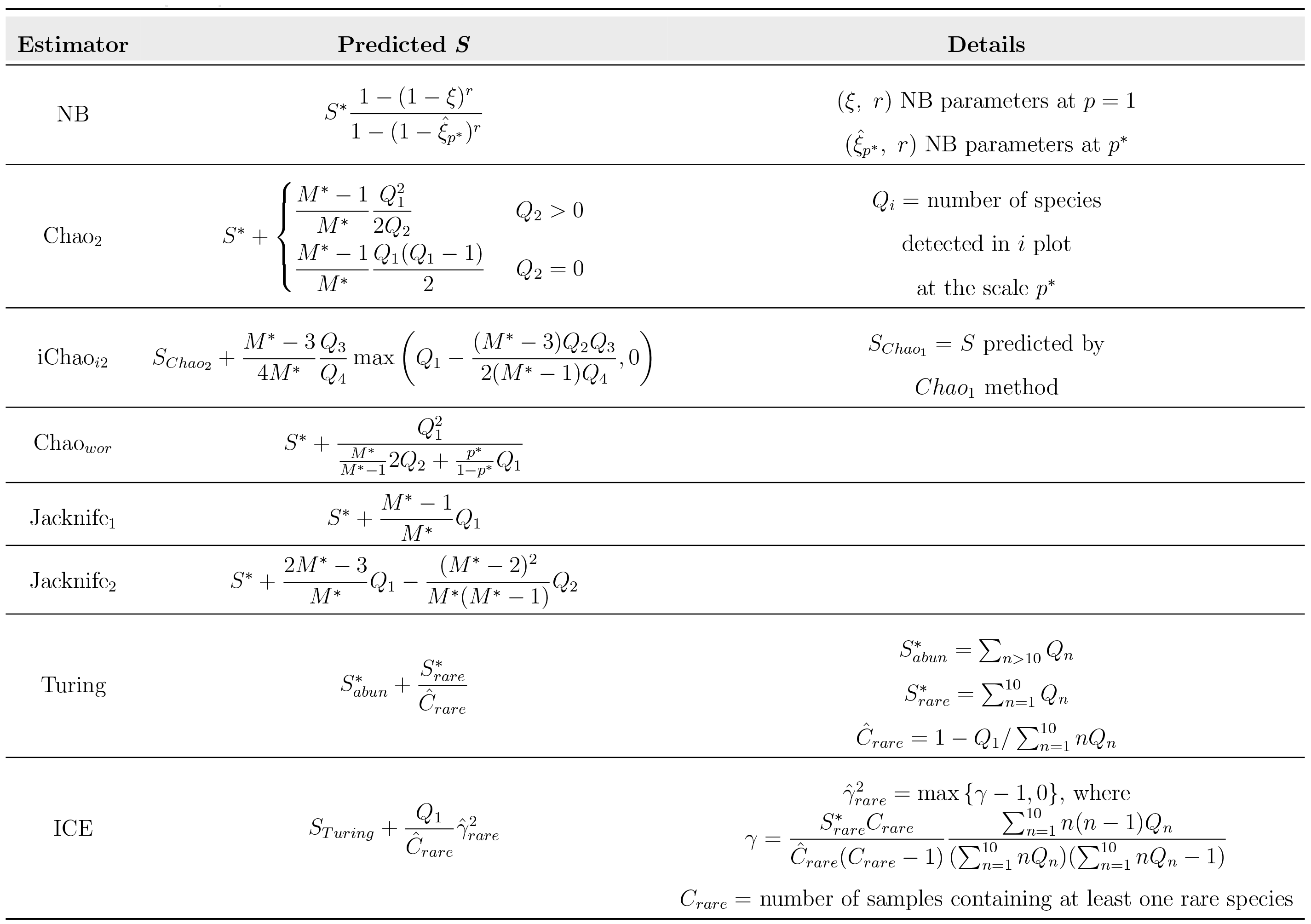
Summary table of the most popular biodiversity estimators for presence/absence data. In formulas, *M*^*^ is the total number of sampled cells. See [30, 11] for more details about non-parametric methods.

Moreover, both parametric and non-parametric methods proposed in the literature do not give any insights on the species abundance at both local or larger scales. Indeed the problem of relating occupancy data with information on species abundance is a relevant issue in theoretical ecology [32, 33, 34]. In particular, given the information on the presence or absence of a species in different scattered plots, one would like to infer its population size or, more generally, the RSA distribution of the forest.

In this paper, we present a general analytical framework to extrapolate species richness and other relevant biodiversity patterns (e.g. RSA, SAC) at the whole forest scale from local information on species presence/absence. Our framework exploits the *form-invariance* property of the Negative Binomial (NB) distribution. Such a distribution emerges as the long time behavior distribution of a birth and death stochastic dynamics, accounting for effective immigration and/or intraspecific interactions [35, 24, 29]. Crucially, the functional form of a negative binomial does not change when sampling different fractions of areas. This property allows for an analytical expression for how parameters of the distribution change across scales. Form-invariance under different sampling efforts is at the core of our approach, however our method can be applied any time the dependence of the distribution on the size of the sampled area can be calculated exactly. We will find an analytical relation between the NB RSA at a given spatial scale and the SAC. Thanks to this function, starting from the empirical SAC constructed at the sample scale from the local presence-absence data (see Eq. (13)), we will be able to:

1. infer species richness at larger scales, thus the SAC up to the whole forest scale;
2. obtain information on species abundances in order to construct the RSA at both local and global scales;
3. introduce and infer the Relative Species Occupancy (RSO), i.e. the distribution of the occurrences (number of occupied cells) across species, at both local and global scales. This biodiversity pattern is a prediction of our modelling framework, can be measured empirically and may be of ecological relevance as it proxies the distribution of species ranges (the area where a particular species can be found) in the ecosystem;

We tested our framework on *in-silico* generated forests and on the two well-studied tropical forests of Barro Colorado Island and Pasoh. We finally compared the global estimates with the abundance-based method proposed in [29].

Before illustrating the details of our approach, we want to highlight differences and similarities between the present work and [29]. Both papers are based on the form-invariance property of the Negative Binomial distribution but, instead of using population estimates at local scales [29], here we require only the knowledge of species’ occurrences at multiple local scales. In other words, the loss of information at one local scale (i.e. for each sample we know if a species is present, but not the number of its individuals) is balanced by the presence-absence information on multiple local scales. Such a generalization of [29] is useful when empirical datasets provide information only on the presence/absence of species. We will show that this will be enough to infer population’s distribution as well.

## 1. Materials and Methods

### 1.1. Theoretical Framework

We denote as *P*(*n*|1) the relative species abundance, – i.e. the probability that a species has exactly *n* individuals – at the whole forest scale (here 1 refers to the whole forest). Note that *P*(*n*|1) should be defined only for *n* ≥ 1, because *S* is the total number of species actually present in the forest, assuming that the area was exhaustively surveyed with no missing species.

Here the RSA at the scale *p* = 1, is postulated to be proportional to a Negative Binomial distribution (NB) [36, 37, 29], 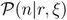 with parameters *r* > 0 and 0 ≤ *ξ* < 1:

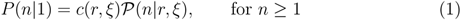

with

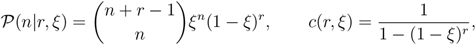

where *c*(*r*, *ξ*) is the normalisation constant. Notice that since *n* ≥ 1, the sum 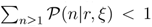 and that is why we need a normalizing factor, taking into account only species with non-zero abundance, which is different from the usual NB normalization. It may be worth to mention here that classically, for a NB distribution, one has *r* ∈ ℕ whereas in our framework *r* ∈ ℝ^+^. Such a distribution can be derived as the steady-state RSA of a simple birth and death stochastic dynamics [36, 37, 29], where *r*, known as the clustering coefficient, models the effects due to immigration events and/or intraspecific interactions, and *ξ* is the ratio between the birth and death rate of a species.

Let us now consider a sub-sample of area *a* of the whole forest and define *p* = *a/A* the sample scale. Assuming that the local RSA is not affected by spatial correlations and/or strong environmental gradients, the conditional probability that a species has *k* individuals in the smaller area *a* = *pA*, given that it has total abundance *n* in the whole forest of area *A* is given by the binomial distribution

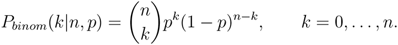

It is worth highlighting that this is where we use the ‘well-mixed’ (or mean-field) hypothesis. This assumption can be tested in the data by looking at the beta-diversity and RSA patterns. If correlation lengths are of the same scale as the system linear size, and the RSA of sub-samples displays the same functional shape, then we can assume that no strong spatial constraints affect the abundance species distribution.

With this information in hand, it can be proved (see [29], Supplementary Materials) that, under the hypothesis that the RSA has a negative binomial form, the RSA at scale *p*, *P*(*k*|*p*), is again proportional to a negative binomial, for *k* ≥ 1, with rescaled parameter *ξ_p_* and the same *r*:

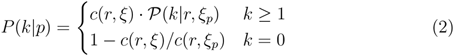

with

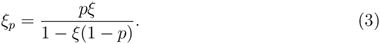

A RSA with the property of having the same functional form at different scales is said to be *form-invariant*.

The *form-invariant* property allows for simple formula describing how birth and death ratios at two different spatial scales are related. Indeed, given the parameters *r* and *ξ*_*p*\*_ of the RSA at the sampling scale *p*\*, we can get the value of *ξ* by inverting (3):

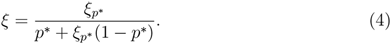

Using (3) to eliminate *ξ* from the last equation, one gets the following relation for the parameter *ξ* at the two scales *p* and *p*\*

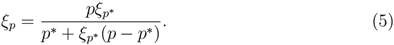

Let now determine the relation between the total number of species at the whole scale *p* = 1, *S*, and the total number of species surveyed at a local scale *p, S_p_*. For the sampling scale *p*\*, in the following, we will use the notation *S*\* ≡ *S*_*p*\*_. Note that, denoting with *S*\*(*k*) the number of species having *k* individuals at the scale *p*\*, one can estimate *P*(*k* = 0|*p*\*) and *P*(*k*|*p*\*) as follows

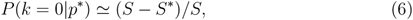

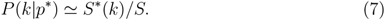

Thus, the total number of species in the whole forest, in terms of the data on the surveyed sub-plot is given by

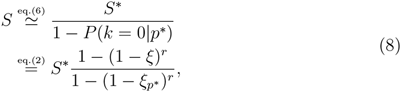

where *ξ* is given by (4).

In general, given two scales *p* and *p*\*, one has the following relation between the number of species at the scale *p*, the one at *p*\* and the RSA parameters (*r, ξ*_*p*\*_) at the scale *p*\*:

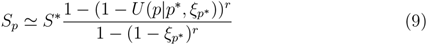

where

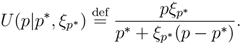

The proposed method is, under the ‘well mixed’ hypothesis, general and not limited to tropical forests.

In the sequel we illustrate how, within the theoretical framework developed so far, it is possible to use presence-absence information on various samples to infer *r, ξ* and then *S*. To start with, let us suppose we surveyed *M*\* cells of the same area *a*. This assumption is not essential to our approach to species estimation at the global scale. It simplifies computations and its implementation (see also next subsection) but can be removed. Suppose we have presence-absence information on each of *M*\* cells. This implies we know *S_p_k__* (i.e. the number of species at scale *p*_*k*_) for *p_k_* = *ka/A, k* = 1,…*M*\*. From eq. (9) we obtain

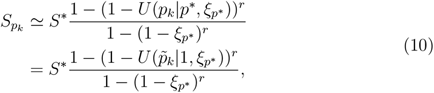

where 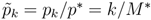 is the fraction of sub-sampled cells. In the last equality of (10) we use that 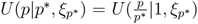 which is obtained from (5) factorizing *p*\*.

The latter equation states that the function of *p* on the righthand side of (9) takes the value *S*_*pk*_ at *p*_*k*_ for *k* = 1,…, *M*\*. For *M*\* ≫ 1, these information allow for a robust estimate of the two unknown parameters *ξ*_*p*\*_ and *r*. Therefore from the empirical values of *S*_*p_k_*_ one can get the parameters *r* and *ξ*_*p*\*_ shaping the RSA at the sample scale *p*\*. From these, one can estimate the *ξ* parameter by using eq. (4) to predict both the number of species at the global scale *S* via (8), the RSA through (1) and the SAC by using (9).

Another important pattern which we can predict with our framework is the relative species occurrence (RSO) distribution, *Q*(*v*|*M*, 1), which gives the probability that a species occupies *v* cells at the global scale, given that the forest can be tiled in *M* equal-sized cells of area *a*. The latter assumption is essential to our derivation of RSO formulae (see eq. (11) and (12) below). Also notice the difference between *M* and *M*\*: in our notation *M* is the number of cells at the global scale whereas *M*\* refers to the fraction *p*\*.

RSO pattern is of ecological relevance as it gives information on the fraction of species that occupy the same amount of area of the ecosystem. For example, if the RSO distribution displayed a small variance unimodal shape, then it means that most of species have similar species ranges. On the other hand if *Q*(*v*|*M*, 1) follows a power law behaviour it indicates a strong heterogeneities in the species ranges.

In order to find an expression for it, we firstly need the probability, *Q*_*occ*_(*v*|*n, M*, 1), that a species occupies *v* over *M* cells at the global scale, given that is has abundance *n*. Under the hypothesis of absence of spatial correlation, this is given by an hyper-geometric distribution. Indeed there are 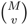 possibilities to choose the *v* filled cells, 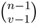 possibilities to distribute *n* species among *v* cells so that no cell is empty, and 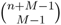 ways to distribute *n* species in *M* cells allowing empty bins. See for example W. Feller, *Introduction to probability theory and its applications*, Chapter 2. Thus we compute

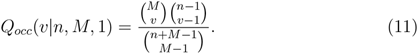

The RSO distribution *Q*(*v*|*M*, 1) can thus be obtained by marginalizing with respect to the abundance *n*:

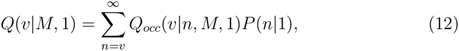

where *P*(*n*|1) is the global RSA given by eq. (1). This series cannot be calculated analytically for arbitrary values of the parameters; nevertheless, it has some regimes which are physically important and can be investigated in more detail. For instance, when *ξ* ≃ 1, *P*(*n*|1) can be approximated by a gamma distribution and, for 0 < *r* < 1 and *v, M* ≫ 1 such that *v*/*M* ≪ 1, one can show that *Q*(*v*|*M*, 1) ∝ *v*^*r*–1^. Since for most forest plots *r* ≪ 1, when *M* and *v* are sufficiently large we expect *Q*(*v*|*M*, 1) = *cv*^−1^, where *c* depends on *M, r* and *ξ*. This prediction is supported by the empirical data we have studied as shown in Figure 3.

**Figure 3:**
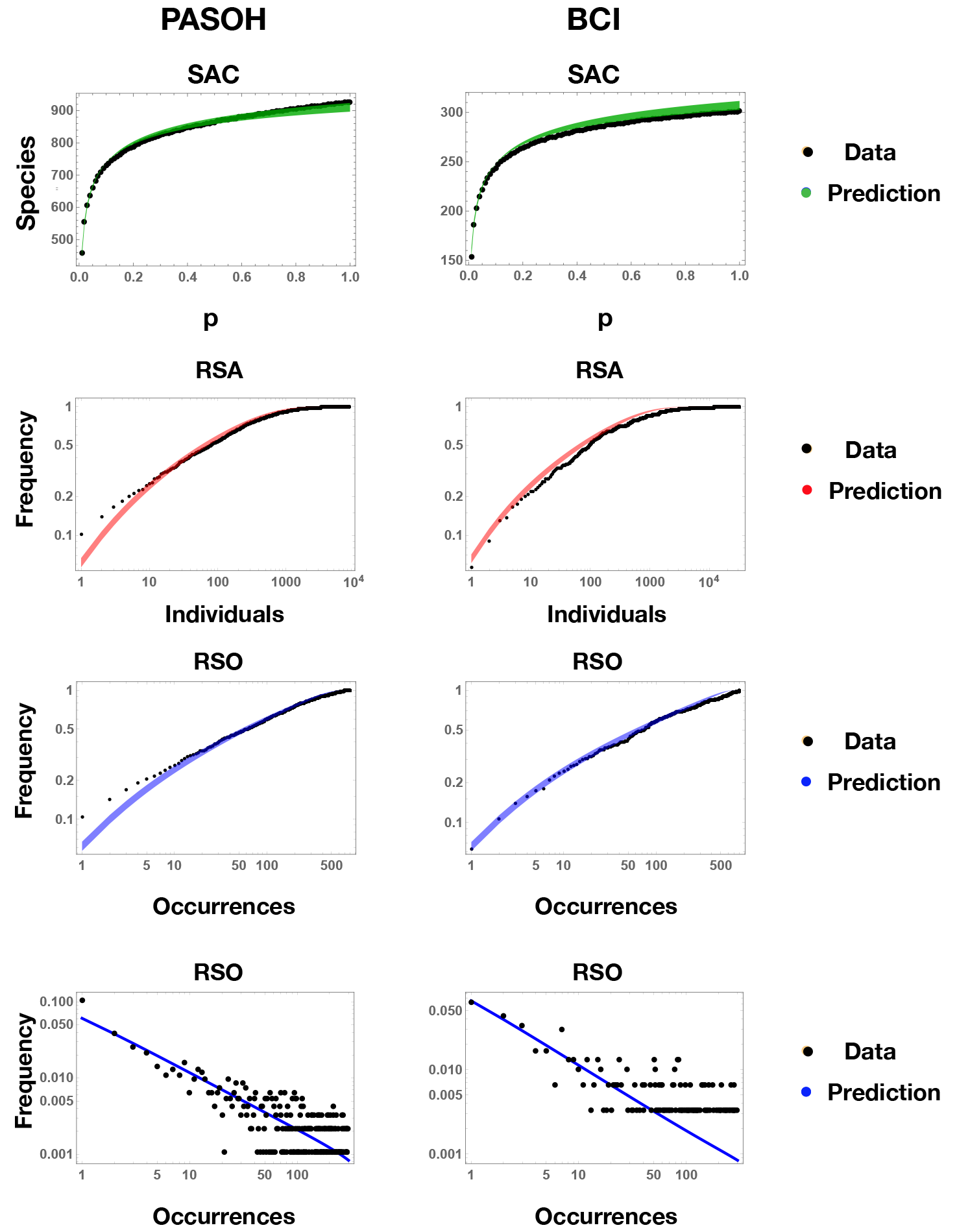
Test on ecological macro-patterns for Pasoh and BCI. For each forest we sub-sample a fraction *p* = 0.1 of the available spatial cells and apply our framework to predict three important ecological pattern at the largest scale at which we have information, *p*\*. In the first row we see the prediction for the SAC curve, which describes how the number of observed species increases with the sampled area, from *p* = 0.1 to 1, corresponding to *p*\* in these tests. In the second row we plot the cumulative empirical RSA, the distribution of abundances across species against the framework prediction in logarithmic scale. Finally, in the third and fourth rows we test the ability of the model to capture the empirical RSO, i.e. the distribution of the occurrences (number of occupied cells) across species in logarithmic scale (third row panel shows the cumulative distribution). In figures, predicted patterns are in the form of confidence intervals obtained from the SAC fitting errors on the *r* and *ξ*_*p*\*_ parameters. For both forests, all the three patterns result to be well described by our framework.

### 1.2. Implementation of the Framework

Our analytical framework consists of the following steps (see Figure 1).

- First, given a set of scattered samples, list the species in it. In formulae, sample *C* = {*c*_1_,…, *c*_*M*\*_}, *M*\* ≥ 2, cells covering a fraction *p*\* of the whole forest in which *S*\* species are observed. To each cell *c*_*i*_, associate a vector 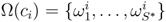, with 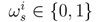, *s* ∈ {1,…, *S*\*}, *i* ∈ {1,…, *M*\*}. The entry 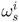 of vector Ω(*c*_*i*_) gives information on the presence/absence of the species s in the cell *c*_*i*_ – i.e. 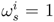 if species *s* is present in cell *c*_*i*_, 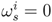 otherwise.
- Compute the empirical species-area curve as follows. From now on, let us suppose that all the *M*\* cells are of equal size *a*. This assumption does not affect the general framework but it simplifies the computation of the SAC. Call *A* the area of the whole forest, so that *p*\* = *M*\**a/A*. At each sub-sampling scale *p*_*k*_ = *ka/A*, with *k* ∈ {1,…, *M*\*}, compute the average number of observed species as

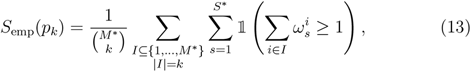

where 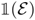 is the indicator function, which equals one when the random event 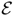 happens and it is zero otherwise. In words: for every scale *p*_*k*_, one should compute the empirical average of the number of the species observed in all subsets of *k* cells. Since computing all subsets of *k* cells among *M*\* is numerically expensive for large *M*\*, in the analyses we computed the average among 100 randomly chosen subsets. Note that computing the species accumulation curve through the empirical average of the number of species in *k* random selected cells, we are neglecting any spatial information. Let us stress once again that null or small spatial correlation is required for a rigorous derivation of our estimates.
- Fit the empirical species accumulation curve with the theoretical equation

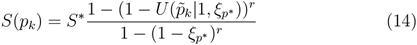

and obtain the parameters (*r, ξ*_*p*\*_) which best describe the empirical curve *S*_emp_(*p*_*k*_). These are the parameters of the NB relative species abundance distribution at the sample scale *p*\*. This protocol allows us to capture some spatial effects in the effective parameters.
- As showed in [29], under the hypotheses of absence of strong spatial correlations due to both inter-specific or intra-specific interactions, strong environmental gradients and abundances distributed according to a negative binomial at the whole forest scale, the RSA distributions at different scales have the same functional form of the RSA at the scale *p*\*, and only the values of the parameter *ξ* changes as a function of the scale. Thus we obtain an analytical form of the upscaled RSA at any scale *p* given we know it at scale *p*\* in term of the equation *ξ*(*p*|*ξ*_*p*\*_) = *U*(*p*|*p*\*, *ξ*_*p*\*_), relating *ξ*_*p*_ = *ξ*(*p*) to *p, p*\* and *ξ*_*p*\*_ = *ξ*(*p*\*). Therefore, using the RSA parameters at scale *p*\* and the upscaling equations (see below), we can predict the total number of species, *S*, at the whole forest scale, *p* = 1.
- The key feature of the method is the possibility, given only presence/absence data, to connect and infer different biodiversity patterns at the global scale. Indeed, we can predict, in addition to the SAC, the RSO, the cell occupancy distribution, and the RSA, the abundance proportions of the *S* species present at *p* = 1.

**Figure 1:**
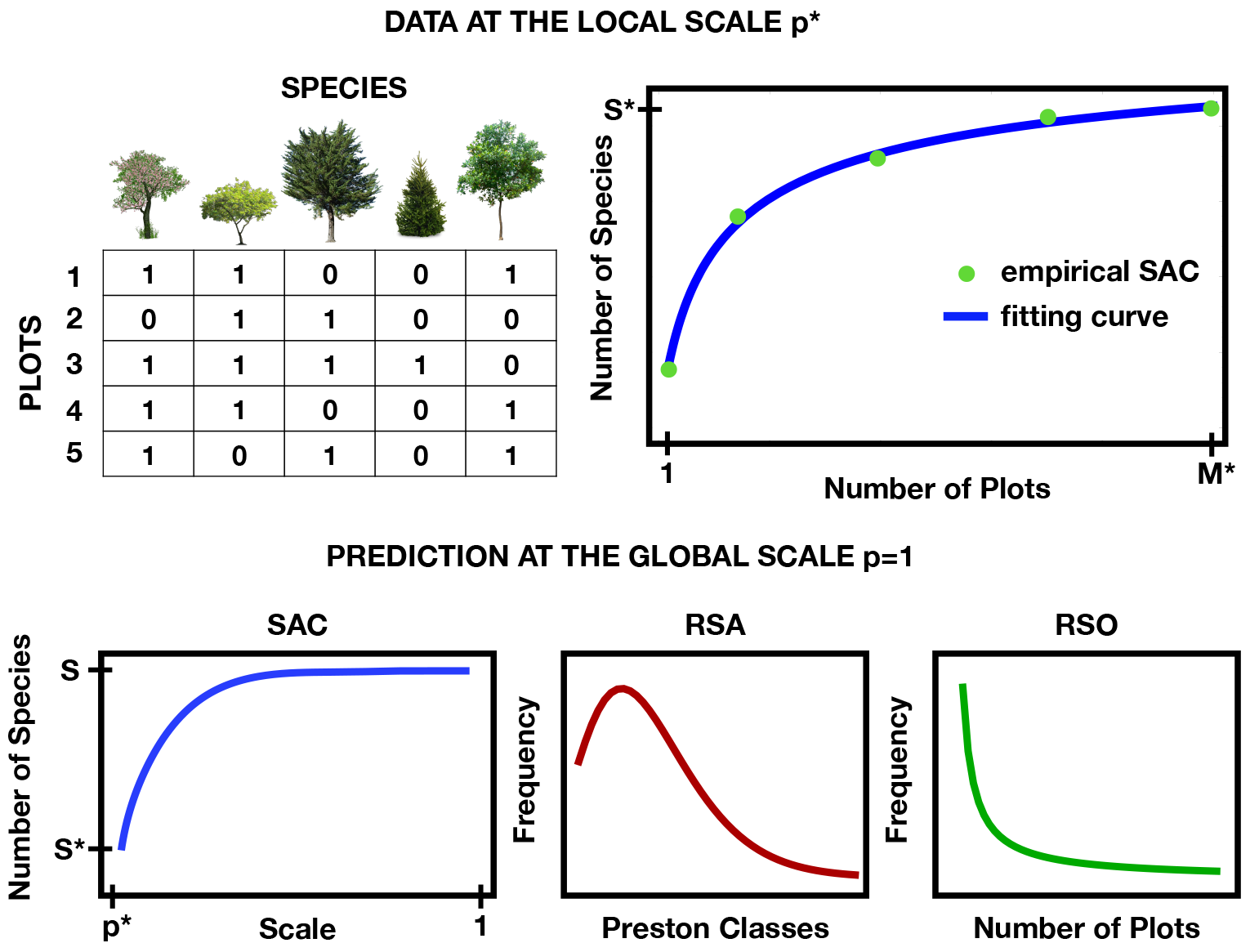
Schematic presentation of our theoretical upscaling framework. It consists of three steps. (A) We start from a dataset in the form of a binary matrix giving information on the presence or absence of *S*\* species within each of the *M*\* surveyed plots. (B) We perform the best fit of the empirically SAC computed via (13). (C) Using the best-fit parameters obtained in (B) and using our upscaling Eqs. (8), (9) and (12), we predict the species richness *S* of the whole forest and three important macro-ecological patterns: the SAC, the RSA and the RSO.

**Figure 2:**
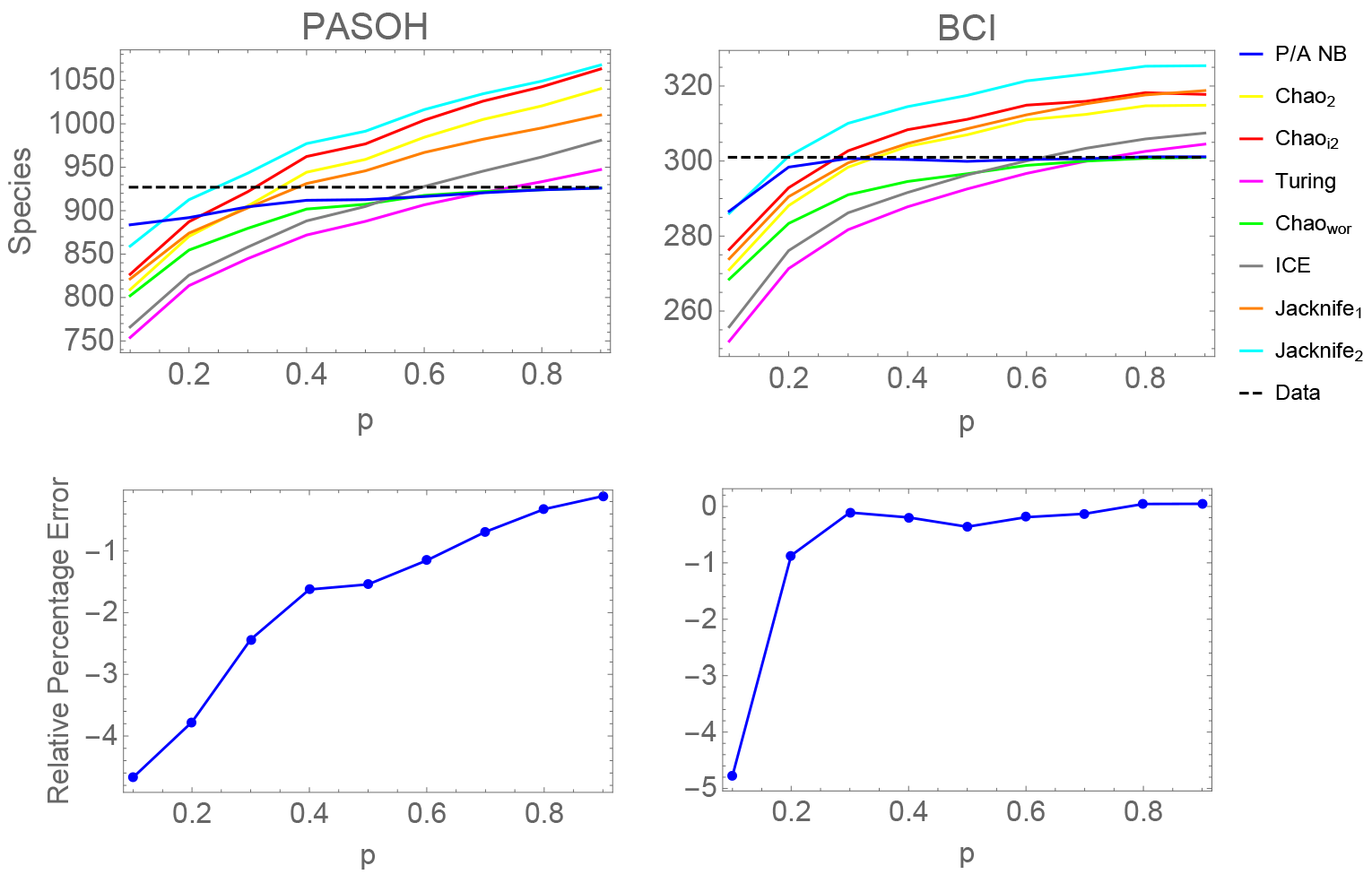
Test from different scales for Pasoh and BCI. For each forest we sub-sample a fraction *p* of *p*\* of the available spatial cells and apply different popular upscaling methods based on presence/absence data (see Table 2) and our method to predict the true number of species, *S*\* (dashed line), observed in our data. While our (P/A NB) and Chao_*wor*_ methods do converge at *S*_*p*\*_ as *p* goes to *p*\*, all the others have a monotonically increasing behaviour due to the independence, in their predictions, of the scale *p*\*. We can see that for both rainforests, our method outperforms all the others. Bottom panels show the relative percentage error (*S*_pred_ – *S*\*)/*S*\* · 100 obtained with our framework between the predicted number of species *S*_pred_ and *S*\*. We find that the method underestimated the true number of species of at most 5%. The larger the sample area, the smaller the relative error.

## 2. Results

### 2.1. Tests on in-silico databases

We test our presence/absence upscaling method on four computer generated forests without and with spatial correlations. Indeed, we expect that in the first case our framework will give more accurate estimates, and we wish to test how the introduction of correlations affect the reliability of our results.

As RSA we choose a negative binomial (NB forest) of parameters *r* = 0.8 and *ξ* = 0.999 and a log-normal (LN forest) with parameters *μ* = 5 and *σ* = 1. Once generated the abundance of every species (*S* = 4974 for the NB forest and 5000 for the LN forest), we distribute the individuals within the forest area, here set equal to a square of 4900 × 4900 units, according to two different processes: at random or according to a modified Thomas process [38, 29, 39] with a clustering radius of 15 units.

We then divide each forest generated as described above into *M* = 98 × 98 units cells and compute the *M* × *S* presence/absence matrix, thus forgetting the information about the species distribution. Finally, we sub-sample the 5% of the cells (corresponding to a fraction *p* = 0.05 of the total forest area) and apply our method to infer the total number of species in each of the four *in-silico* forests. We also compared our results to those obtained by accounting for the data on species populations with an abundance-based upscaling framework developed and tested in [29]. In the case of the NB forest, the two methods performed very well for both the random and the clumped distribution (i.e. individuals distributed on the space according to a Thomas point process) with an average prediction error below 1% in absolute value (see Table 1). In the Thomas distributed forests, the error increased, although remaining around 3% for the presence/absence method and around 7.5% for the abundance-based one (using maximum likelihood methods. The latter percentage error can be improved using calibrated statistical method for the single fit). Thus, with respect to the degree of individuals’ clustering, the new framework seems to give more robust estimates than the second one. This is due to two main reasons: 1) For the presence-absence case, we fit the empirical SAC, which has a very smooth functional shape, and it is easy to describe through our analytical SAC. On the other hand, the RSA displays a more complex and variable shape and thus fitting it with the NB is a more delicate task (indeed we find sensible differences on the accuracy using different statistical methods for the fit); 2) Binary data on which the empirical SAC is based are less sensitive to sampling fluctuations.

**Table 1:** Predictive error for three generated forest (characterized by a log-normal and a negative binomial RSA) having individuals distributed according to a high clustering Thomas process and at random. Tests were performed by sampling a fraction *p* = 0.05 of each forest and by applying our framework (P/A columns) and the abundance-based method (RSA columns) to predict the true number of species *S* (5000 for the LN forest and 4974 for the NB forest). For each estimated *S*_pred_, the average relative percentage error (*S*_pred_ – *S*)/*S* · 100 between the true number of species and the predicted one is shown together with the corresponding standard deviation. Results are relative to 100 iterations.

| Forest RSA | Spatial Distribution |  |  |  |
| --- | --- | --- | --- | --- |
|  | Random |  | Thomas |  |
|  | P/A | RSA | P/A | RSA |
| Log-normal | $3.1 \pm 0.51$ | $7.6 \pm 0.52$ | $2.5 \pm 1.8$ | $7.2 \pm 3.1$ |
| Negative binomial | $-0.50 \pm 0.34$ | $-0.52 \pm 0.28$ | $-0.81 \pm 1.6$ | $-0.60 \pm 1.7$ |

### 2.2. Tests on real databases

We finally test our method on sub-samples taken from two empirical forest data for which we have informations on both species occurrence and abundances. In particular we extract abundances of tree species observed in 50ha of rainforests from Pasoh (Malaysia) and Barro Colorado Island (Panama) together with the spatial locations of each of their individual.

Firstly, we divide both forest data into a grid consisting of *M* = 800 equal-sized cells of area 625 m^2^ and we derive the *M* × *S*\* presence/absence matrix for the *S*\* observed species (*S*\* = 927 for Pasoh forest and 301 for BCI). We then sub-sample species occurrence for different fractions 0 < *p* < *p*\* of the cells and apply our framework to infer the number of species and other biodiversity patterns (RSA, RSO and SAC) at the corresponding largest empirically-observable scale *p*\*, for which we know the ground truth.

We compared our results on species richness obtained only from presence-absence data with the most popular non-parametric indicators proposed in the literature [30, 11], which are summarized in Table 2. We found that our method outperforms all the others for both BCI and Pasoh forests. We also remark that all these methods have the further limitation that they can only infer the total species richness, without allowing for an estimate of the abundances’ and occurrences’ distributions, i.e the shape of the RSA and the RSO.

Indeed, as shown in Figure 3, from the local presence-absence data, we can reconstruct, among the SAC, the RSA at the whole tropical forest scale, thus relating species occurrence data with information on the abundances. In particular we can see that the inferred RSA are statistically comparable with the empirical ones obtained by using all the information on species’ abundances which we deleted before applying our method.

Another biodiversity pattern that we can infer from our framework is the RSO. As shown in Figure 3, we find that, as for the RSA, this pattern seems to have a universal form which can be well described and correctly inferred through our neutral approach. Also, our finding suggests that, when spatial effects are negligible, the RSO distribution has a wide range of values in which it is well approximated by a universal power law, regardless of the details of the populations’ dynamics. One may assume that this latter is driven by a simple stochastic process with constant per capita birth and death rates. Such a slow decay of *Q*(*v*|*M*, 1) indicates that species in real systems exhibit huge variations in their occurrences, which may be weakly correlated to species’ habitat preferences or environmental heterogeneities. We should expect strong asymmetries among their occurrences: for instance, if we tile up a landscape into *M* = 1,000 elementary cells, then about a third of all species should live in less than 1% of them; whereas about 1.5% of the species should be found in more than 90% of the total cells (see Figure 3).

We highlight that the SAC (green line), the cumulative RSA (red line) and cumulative RSO (blue line) predicted patterns in Figure 3 have not been obtained through the fit of some parameters, but they have been analytically predicted through our upscaling equations (1), (12) and (9). The only fitting occurs at the scale *p* = 0.1*p*\* using the empirical SAC to parametrize eq. (13). In other words, by fitting species occurrence data at the sample scale, our framework allows to estimate: 1) The RSA at the sample scale; 2) The SAC, the RSA and the RSO at larger scales. We provide an open source R code that performs the above estimates giving as input only the presence-absence matrix data.

After testing our model on controlled computer generated data and real forest sub-samples, we apply our framework to predict the species richness of the two tropical forests. Moreover, we compare our results to those obtained with the upscaling framework based on RSA pattern previously developed and tested in [29] by our group.

We therefore predict, through the presence/absence method, the species richness at the whole forest scale (*p* = 1) for BCI and Pasoh tropical forests. Figure 4) shows the prediction of the overall (and unknown) SAC for a scale ranging from 50 to 14000 hectares for the Pasoh (*p*\* ≈ 0.0036) and to 1560 for the BCI (*p*\* ≈ 0.032). The blue curves represent the prediction obtained only using presence-absence data whereas red curves are the SAC inferred by exploiting also the information about species’ population through the abundance-based method (see [29]).

**Figure 4:**
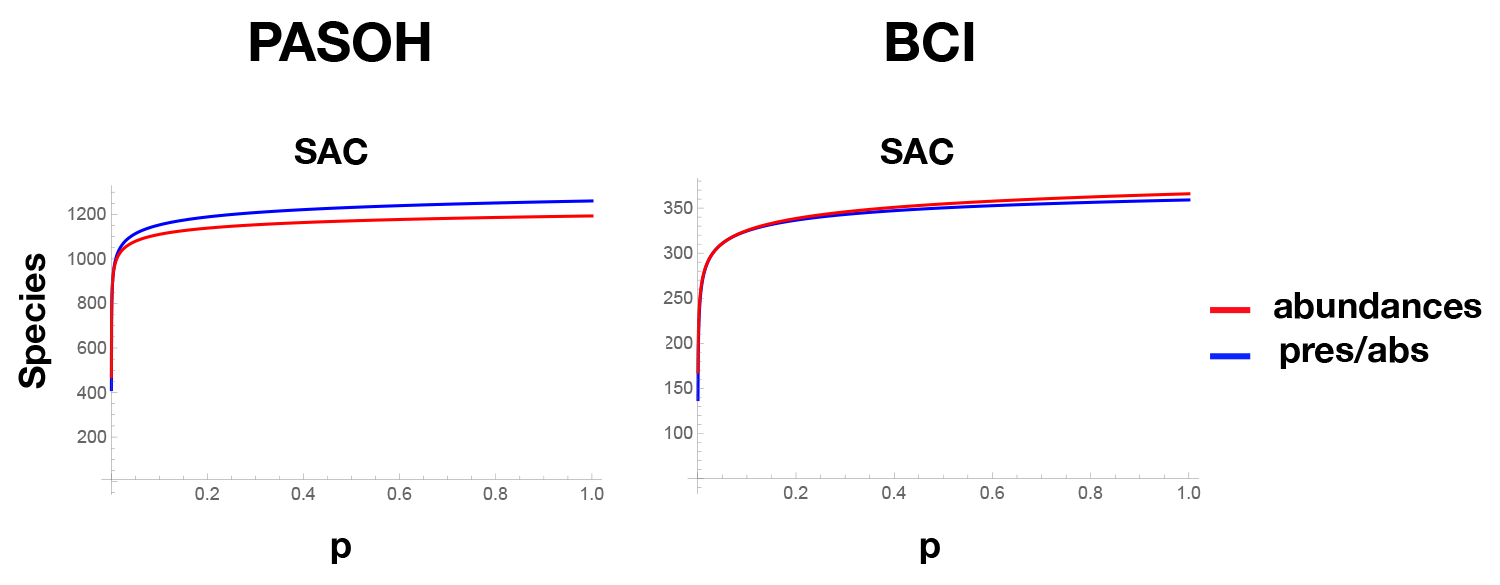
SAC predicted for Pasoh and BCI using abundance method versus presence/absence method. Using all the available data for both tropical forests, we compare the prediction for the SAC curve obtained by the abundance method [29] with the results obtained with the presence/absence framework presented here. At the whole forests’ scale *p* = 1, the two predictions are 3*σ* compatible 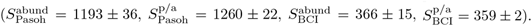

We find that the two methods give comparable results for both the databases, a confirm of the robustness of the theoretical framework.

## 3. Discussion

In this work we proposed and tested a novel rigorous statistical framework to upscale ecological biodiversity patterns from local information on species occurrence data. Different upscaling approaches have been proposed in ecological literature [26, 28, 40, 4, 9, 10, 11, 18, 41, 40]. However, to the best of our knowledge, they have not been generalized to the case of binary data. The present paper provides a generalization of the method recently proposed in [29] to presence-absence information. In [29] species abundance distributions at one given scale was required to make predictions at global scale, whereas the present approach allows to extract abundance distributions at any scale from species occurrence data in multiple small scale samples. The underlying hypotheses that we need in order to perform these estimates is that the RSA at a given scale is a negative binomial distribution, a RSA that arises naturally as the steady-state species abundance distributions for ecosystems undergoing simple birth and death dynamics [42, 24]. The negative binomial is a simple and versatile distribution that depending on its parameters can display an interior mode or log-series like behaviour, i.e. it can accommodate different RSA shapes. Therefore we can use the same RSA function to reproduce different ecosystems’ RSA, as those typically observed in real ecosystems [43, 44, 45, 35, 46, 47, 48, 49, 24]. Even more generally, by using mixtures of negative binomials - a case for which our framework still works - we can fit more complex RSA shapes (see [29]).

Furthermore, we introduce a new descriptor/measure of biodiversity within an ecological community, the RSO, which describes the distribution of species occurrences in scattered plots. The RSO distribution displays a fat tail, indicating that many species typically occupies only few scattered plots, while only very few species are pervasive and are found in most of the plot. Our prediction is that this property is not particular for the dataset here considered, rather it is another emergent patterns [50, 24] pervasive in highly biodiverse ecosystems. Our framework gives directly all parameters of the RSO by solely fitting the SAC curve, through which one can obtain the *r* and *ξ* parameters, which well describe both the RSA and the RSO distributions at all spatial scales of interest.

Expanding the ability to upscale species richness and obtain abundance distributions from presence-absence data is of fundamental importance in many contexts, where abundance information are not available or trustable. This is particularly true for microbial or marine (e.g. plankton) ecological data obtained from metagenomics [51] and 16S ribosomal gene sequences [52]. The use of sequence-based taxonomic classification of environmental microbes has exploded in recent years [52, 53, 51, 31] and these approaches are becoming a standard method for characterizing the biodiversity of both prokaryotes and eukaryotes [53]. Thanks to advance in high throughput sequencing we begin to be able quantifying the vast number of microbes in our environments, expanding our knowledge on microbial diversity [31]. However, large fractions of the sequence reads remain unclassified [51] and also species abundance estimated have a very high uncertainty [31]. Thus, being able to estimated species richness and abundance distributions from species occurrence data may lead to a big step-forward in the taxonomic classification of microbial ecosystems.

To summarize, this flexible analytical method provides, from local presence/absence information, robust estimates of species richness and important macro-ecological patterns of biodiversity (SAC, RSA, RSO), as tested in both *in-silico* generated and two rainforests. The method may be applied to any database in the form of a binary matrix, where presence/absence features (tree species in our case) are detected across different samples.

## Acknowledgments

This preprint has been reviewed and recommended by Peer Community In Ecology (https://dx.doi.org/10.24072/pci.ecology.100009).

## Data Availability

All data are publicly available. The Pasoh and Barro Colorado Island datasets are provided by the Center of Tropical Research Science of the Smithsonian Tropical Research Institute (https://stri.si.edu/). R codes are available at https://github.com/annatovo/Inferringmacro-ecological-patterns-from-local-species-occurrences.

